# Extracting dynamical understanding from neural-mass models of mouse cortex

**DOI:** 10.1101/2021.12.22.473927

**Authors:** Pok Him Siu, Eli Müller, Valerio Zerbi, Kevin Aquino, Ben D. Fulcher

## Abstract

New brain atlases with high spatial resolution and whole-brain coverage have rapidly advanced our knowledge of the brain’s neural architecture, including the systematic variation of excitatory and inhibitory cell densities across the mammalian cortex. But understanding how the brain’s microscale physiology shapes brain dynamics at the macroscale has remained a challenge. While physiologically based mathematical models of brain dynamics are well placed to bridge this explanatory gap, their complexity can form a barrier to providing clear mechanistic interpretation of the dynamics they generate. In this work we develop a neural-mass model of the mouse cortex and show how bifurcation diagrams, which capture local dynamical responses to inputs and their variation across brain regions, can be used to understand the resulting whole-brain dynamics. We show that strong fits to resting-state functional magnetic resonance imaging (fMRI) data can be found in surprisingly simple dynamical regimes—including where all brain regions are confined to a stable fixed point—in which regions are able to respond strongly to variations in their inputs, consistent with direct structural connections providing a strong constraint on functional connectivity in the anesthetized mouse. We also use bifurcation diagrams to show how perturbations to local excitatory and inhibitory coupling strengths across the cortex, constrained by cell-density data, provide spatially dependent constraints on resulting cortical activity, and support a greater diversity of coincident dynamical regimes. Our work illustrates methods for visualizing and interpreting model performance in terms of underlying dynamical mechanisms, an approach that is crucial for building explanatory and physiologically grounded models of the dynamical principles that underpin large-scale brain activity.

## Introduction

Recent advances in neuroimaging have produced intricate maps revealing the complexity of the brain’s microscale circuits, with whole-brain coverage. Analyzing and integrating these data have uncovered new patterns of brain organization, including the systematic spatial variation of gene expression (1, 2), cytoarchitecture (3), neuron densities (4), cortical thickness (5), axonal connectivity (6), cognitive function (7), and local dynamical properties (8) like intrinsic timescales (9, 10). Existing evidence suggests that, to a good first approximation, these properties vary together along a dominant hierarchical axis in mouse and human (1, 2, 11).

To understand the functional role of observed physiological patterns, like systematic spatial variations in brain architecture, we need a way of simulating their effect on wholebrain dynamics. Physiologically based brain models achieve this, using methods from statistical physics to capture the dynamics of large populations of neurons and their interactions (12, 13). Neural population models can capture the complex spatiotemporal dynamics in modern neuroimaging datasets, including persistent activity, intermittent oscillations, and multi-stability (14), and have successfully reproduced a wide range of experimental phenomena, from the alpha rhythm to seizure dynamics (13). The physiological formulation of these models means that their variables and parameters encode interpretable and biologically measurable properties of neural circuits, like the strengths and timescales of interactions between neuronal populations. This allows them to provide a unique mechanistic account of whole-brain dynamics that can be validated against both physiological experiments and the dynamical patterns observed in neuroimaging experiments.

While most existing brain models involve dynamical rules that are spatially uniform (e.g., the same model parameters in all brain areas), recent work has begun to investigate the effect of non-uniform dynamical rules, constrained by emerging brain-atlas datasets. An early example is the work of Chaudhuri et al. (15), which incorporated a variation in recurrent excitation corresponding to that of measured spine count in the macaque. More recent work in human has incorporated spatial heterogeneity in model parameters with: the MRI-derived T1w:T2w map (16); T1w:T2w, the first principal component of gene transcription, and an inferred excitation:inhibition ratio (17); a linear combination of T1w:T2w and the principal resting-state functional connectivity (FC) gradient (18); and a fitted parametric variation that recapitulated an interpretable hierarchical variation (19). These papers have reported improved out-of-sample model fits to empirical data, evaluated according to a range of summary statistics of the resulting dynamics (most typically FC), and provided insights into how spatial variation in biological mechanisms (like recurrent excitation) may underpin whole-brain dynamical regimes. While these studies demonstrate the promise of producing more accurate predictions of measured brain dynamics by incorporating regional heterogeneity—constraining to physiological data, or through large-scale parameter fitting (19)—the resulting models are correspondingly complex and challenging to interpret in terms of the mechanisms which underpin their dynamics. For example, the tools of dynamical systems have the potential to reveal the dynamical features that improve model fits to data, including the bifurcation structure that defines the accessible dynamical regimes and the range of such regimes that different brain areas can access, including their vicinity to critical points (16, 19–22). In this work, we show that analyzing the dynamical response of individual brain regions to inputs using bifurcation diagrams provides an understanding of model behavior in terms of accessible dynamical regimes, an approach that is particularly valuable for understanding the increased complexity of spatially non-uniform models.

The mouse is an ideal organism to develop comprehensively constrained physiologically based models of brain dynamics, but models of the mouse brain have been relative few (23–25) compared to the large number of studies of human cortex. Compared to human, there is an abundance of high-resolution, whole-brain physiological data in mouse (2), including directed tract-tracing axonal connectivity data (6, 26), high-resolution gene-expression maps (27), and celldensity atlases (4, 28). High-quality whole-brain neuroimaging data using fMRI in mouse is also available, allowing us to evaluate model predictions in the resting state (29, 30) and as a result of targeted manipulations (31, 32). Prior work has shown that FC is strongly constrained by direct structural pathways (33), and prior dynamical models have reported the ability of coupled dynamical models to reproduce FC structure, especially when modeling using matching individual structural connectivity (24). In this work, we develop a neural-mass model of mouse cortical dynamics, and aim to understand the dynamical regimes in which it best captures resting-state fMRI data in mouse. We also aim to characterize the impact of incorporating spatial variations in excitatory and inhibitory cell densities as spatial variations in model parameters from a dynamical systems perspective.

## Methods

As illustrated in Fig. 1, we developed a neural mass model of 37 mouse cortical areas, comprising a simple Wilson–Cowan local dynamical model (Fig. 1A) coupled via a directed structural connectome (Fig. 1B). These regions are shown on the mouse brain in Fig. 1C, colored by their relative excitatory cell densities. Of the 38 cortical regions reported in Oh et al. (6), we excluded the frontal pole (FRP) due to its small size (likely contributing to noisy, outlying values of excitatory and inhibitory cell densities (4)). In visualizing our results, we grouped cortical regions according to six functional labels: Somatomotor, Medial, Temporal, Visual, Anterolateral, and Prefrontal (26) (see Table S1 for full list).

**Fig. 1.**
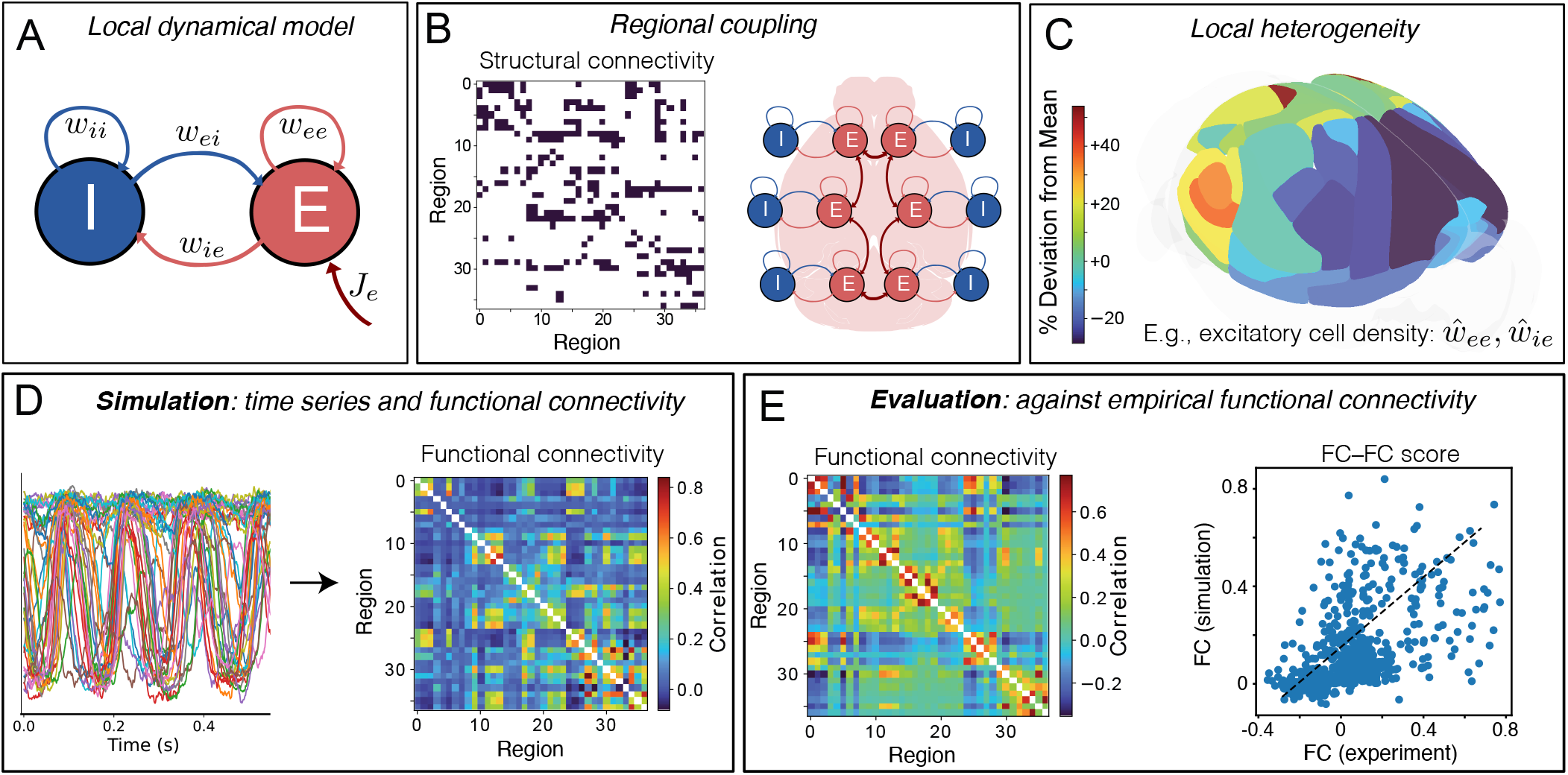
Simulating and evaluating a coupled neural-mass model of mouse cortical dynamics. **A** The dynamics of individual brain regions follow the Wilson–Cowan equations (34, 35) which govern interactions between local excitatory (*E*) and inhibitory (*I*) neural populations. **B** Regions are coupled together by connections defined by the AMBCA (6), represented as a directed adjacency matrix (connections shown black). A schematic shows how these long-range structural connections couple local cortical regions via excitatory projections (13). **C** Heterogeneity in local model parameters can be introduced as a perturbation that follows the measured variation in excitatory and inhibitory neural densities. Here the variation in excitatory cell density is plotted across the 37 mouse cortical areas as deviations relative to the mean level (green), using brainrender (37) and data from Erö et al. (4). **D** Model simulation yields activity time series for each brain region, from which pairwise linear correlations (functional connectivity, FC) are computed. **E** Model simulations are evaluated against empirical FC, averaged across 100 mice, as the Spearman correlation between all unique pairwise FC values, yielding an FC–FC score, *ρ*_FCFC_.

As shown in Fig. 1A, a given brain region consists of both an excitatory (*E*) and an inhibitory (*I*) neural population, whose dynamics are governed by the Wilson–Cowan equations (34, 35). Brain regions are coupled via long-range excitatory projections using a binary, directed connectome from the Allen Mouse Brain Connectivity Atlas (AMBCA) (6, 36) (Fig. 1B). Simulating the model yields dynamics for the *E* and *I* populations; we take the activity time series of the excitatory population to evaluate the similarity of pairwise linear correlation structure as functional connectivity (FC), shown in Fig. 1D. To assess the goodness of fit, we compare this simulated FC to an empirical FC calculated on a mouse fMRI dataset (Fig. 1E). The goodness of fit is assessed as a Spearman correlation coefficient computed between all pairs of FC values from the empirical data and the model (Fig. 1E). Spearman’s correlation coefficient was used instead of Pearson’s correlation coefficient to capture a potentially nonlinear but monotonic relationship.

As our main aim was to develop tools to understand the distributed dynamics of neural mass models, we favored simplicity in focusing on the Wilson–Cowan (W–C) model relative to alternative models. In addition, its physiological formulation is crucial for mapping to experimental cell-density data, as its parameters encode measurable properties with physical units that can be constrained by such data. The W– C model also exhibits a wide range of dynamical behaviors, including bifurcations, hysteresis, stable fixed-points (attractors), and limit cycles (oscillatory attractors) (34, 35, 38), that are common features of dynamical systems in general, including more complex biophysical neural population models. We use a formulation of the Wilson–Cowan equations based on the mean firing rates of coupled populations of excitatory and inhibitory neurons, as

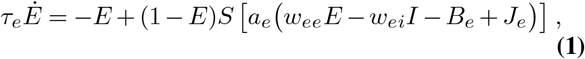

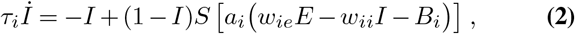

where *E* and *I* are the mean firing rates of the excitatory and inhibitory populations, respectively (Hz); *S*(*v*) = *h/*(1 + exp(^_^*v*)) is the sigmoidal firing-rate function, *h* is the upper bound for the sigmoid function representing a maximal population firing rate (Hz); *a*_*e*_, *a*_*i*_ control the gradient scaling for the sigmoid function (V^_1^); *w*_*xy*_ are the coupling weights from population *y* to population *x*, where *x* and *y* correspond to excitatory (*e*) or inhibitory (*i*) populations (V s); *B*_*e*_, *B*_*i*_ are the firing thresholds for excitatory/inhibitory cells (V); *J*_*e*_ is the voltage induced by external current injected into the excitatory cells, defined below as a weighted sum over external inputs (V); and *τ*_*e*_, *τ*_*i*_ are the time constants of excitation and inhibition respectively (s).

Neural masses, corresponding to cortical areas, were coupled via projections between excitatory populations through the external current term, *J*_e_. For a given region *a*, 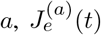 is computed as

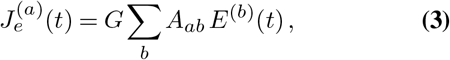

where *G* is a global coupling constant (V s), *A*_*ab*_ is the adjacency matrix corresponding to the structural connectome (unweighted here), and *E*^(*b*)^ is the excitatory activity of region *b* (Hz). It is helpful to define the quantity

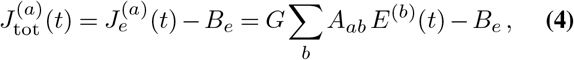

as the total input that includes the constant offset *B*_*e*_. We will use this to understand the dynamical response of a brain area to its net input.

In addition to this ‘homogeneous’ model, in which the parameters are identical for all brain regions, we also analyze a heterogeneous model, in which the coupling parameters, *w*_*ij*_, vary across regions. We calibrate this variation to estimated cell-density data (4), by making the assumption that local connectivity from excitatory and inhibitory cells is uniform, and thus that coupling strengths from a given population are proportional to the density of cells of that population. Thus, we adjust the coupling parameters corresponding to outputs from the excitatory population, *w*_*ee*_ and *w*_*ie*_, according to measured variations in excitatory cell density across cortical areas, and adjust *w*_*ii*_ and *w*_*ei*_ according to measured variations in inhibitory cell density. Defining nominal parameter values as *ŵ*_*xy*_ (for *x* and *y* taking *i* and *e*), we can then define linear parameter perturbations for a given region *a* as:

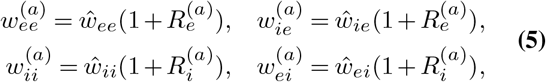

where rescaling factors, *R*_*e*_ and *R*_*i*_, represent relative variations in excitatory and inhibitory cell density, respectively (see Fig. 1C for a visualization of how excitatory cell density varies across cortical areas). To map cell-density measurements to corresponding *R*_*e*_ and *R*_*i*_ values, we first *z*-score normalized raw excitatory and inhibitory cell-density data, as *e*^(*a*)^ and *i*^(*a*)^, respectively, across all regions, *a*. We then defined a simple proportional mapping to model parameters via a single scaling parameter, *σ ≥* 0, as

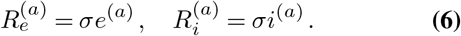

In this formulation, setting *σ* = 0 sets all 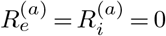 and reproduces the spatially homogeneous model; increasing *σ* increases the level of variation in coupling parameters across areas. Note that there is much scope for defining more complex mappings involving more new parameters, but defining the mapping from cell densities to model parameters in this simple, single-parameter scheme allows us to more clearly tackle our main aim to investigate how the model’s dynamical features are shaped by such variation.

For a given system of coupled ODEs defined above, dynamics were simulated using The Virtual Brain (TVB) (23, 39), yielding simulated time series for each region. The system was driven by white noise with a mean *µ* = 0 and standard deviation *σ* = 1.3 × 10*™*5 using the Euler–Maruyama method with a fixed time step, Δ*t* = 0.1 ms, for a total simulation length of 1.2 × ×105 ms, (or 2 min at 1000 Hz). Initial transients of 1 s (1000 time steps) were removed from all simulations to focus on the model’s steady-state dynamics. Importantly, our aim was to understand the dynamical properties of the model that enable it to match the statistics of measured dynamics, and chose not to adjust the model output, *E*^(*a*)^(*t*), through a simulation of the hemodynamic response function to match the fMRI measurement (but could be done in future using e.g., a convolution of a canonical hemodynamic response (40) or a biophysical model (41, 42)). Models were assessed on their ability to reproduce the pairwise linear correlation structure (functional connectivity, FC) of empirical mouse fMRI data, as the Spearman correlation between predicted and measured FC values: the FC–FC score, *ρ*_FCFC_. While we focused here on reproducing pairwise linear correlations using *ρ*_FCFC_, we note that a more comprehensive evaluation of model fit, incorporating aspects of local dynamics and dynamic FC properties, will be important for future investigations to more fully evaluate the rich patterns contained in the dynamics (17, 43, 44). To account for variability in simulated model dynamics due to a finite simulation time and different random seeds, we computed *ρ*_FCFC_ for 40 repeats of each simulation using different random seeds. Code for reproducing the simulations and analysis presented here is available at https://github.com/Spokhim/MouseBrainModelling.

## Results

Here we aim to understand the dynamical principles underlying coupled dynamical models using a neural mass model of the mouse cortex. First, we investigate the spatially uniform case in which all brain regions are governed by identical dynamical rules. Focusing on model behavior in the vicinity of saddle-node and Hopf bifurcations, we characterize the model’s dynamical regimes that best capture empirical FC structure. We then investigate the spatially heterogeneous case, in which regional variations in parameters are introduced according to variations in excitatory and inhibitory cell-density maps (4), which shape the model’s local bifurcation properties and resulting dynamical regimes.

### What dynamical features drive high model performanceã

We first characterize the model’s dynamical regimes that best capture the pairwise correlation structure of experimental mouse fMRI, with the aim to understand how the positioning of individual nodes (brain regions) around specific types of bifurcations affects the model’s ability to capture empirical FC. In this section we focus on a homogeneous model, in which all brain areas are governed by the same dynamical rules, but differ in their inputs from other regions (via the connectome). We characterize the model’s behavior in each of three regimes: (i) in the vicinity of a single stable equilibrium, which we denote as the ‘Fixed Point’ regime (using parameters adapted from Sanz-Leon et al. (39)); (ii) in the vicinity of a bistable region separated by saddle-node bifurcations, which we denote as the ‘Hysteresis’ regime (using parameters from Heitmann et al. (45)); and (iii) in the vicinity of a pair of Hopf bifurcations, denoted as the ‘Limit Cycle’ regime (using parameters from Borisyuk and Kirillov (46)). Parameter values for each of these three regimes are given in Table S2. Bifurcation diagrams of excitatory firing, *E*, as a function of net external input, *J*_tot_ = *J*_*e*_ − *B*_*e*_, are plotted for the Fixed Point regime (Fig. 2D), Hysteresis regime (Fig. 2E), and Limit Cycle regime (Fig. 2F). These plots show how stable states of *E* vary with *J*_tot_ as solid lines (with unstable states shown for the bistable regime in Fig. 2E and lower and upper limits of a limit-cycle oscillation in Fig. 2F). They thus capture the rules underlying the dynamical behavior of individual brain regions in response to their aggregate input from other brain regions, *J*_tot_, with each parameter setting defining a qualitatively different set of accessible dynamics, and types of response to inter-regional inputs. Importantly, these basic bifurcation structures, and the insights we gain from them, are not specific just to the W–C model, but are common features of many dynamical models (47).

**Fig. 2.**
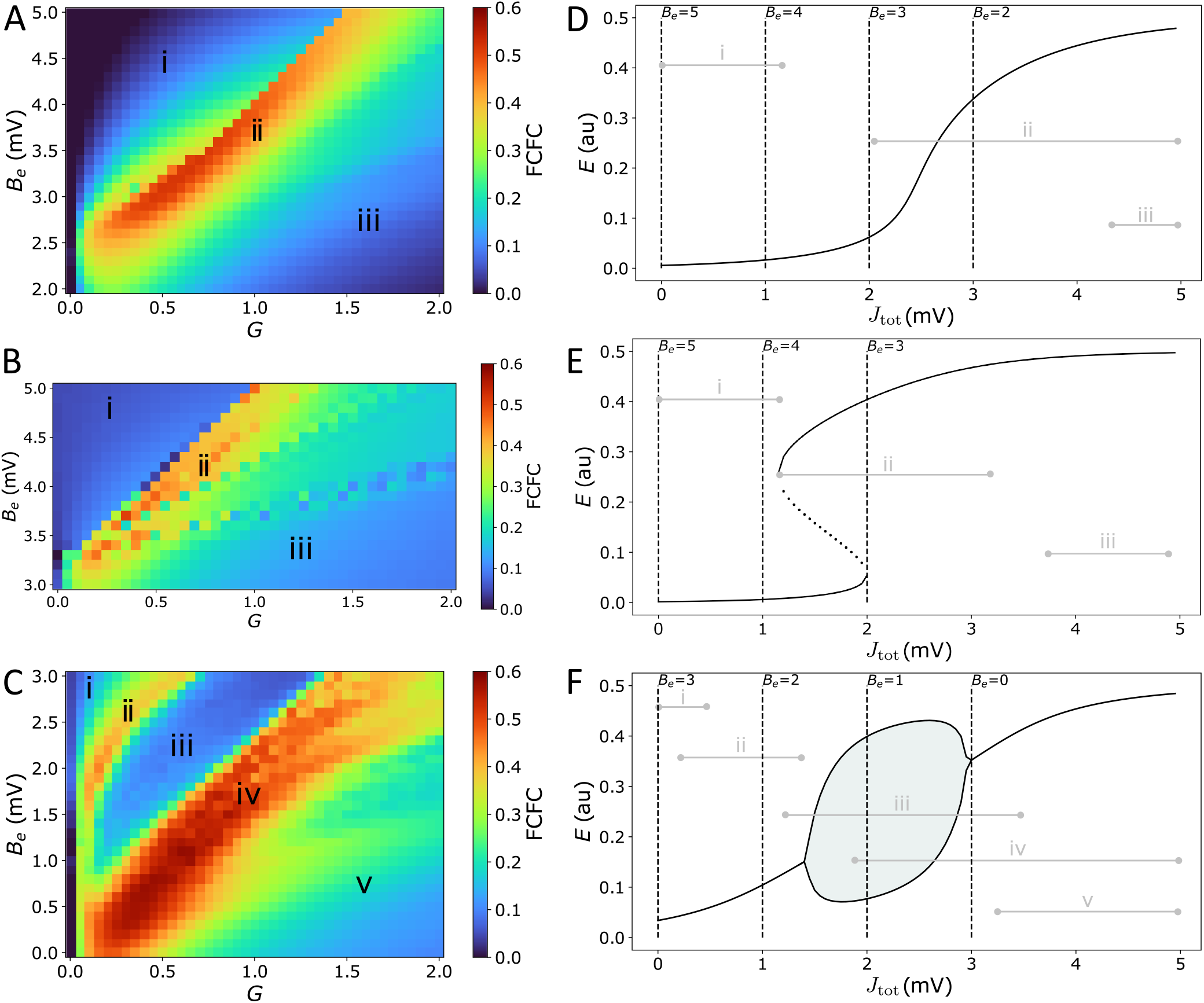
Model performance is highly sensitive to the types of dynamical features available to the coupled dynamical network, with high FC–FC found near bifurcations and where external inputs have strong dynamical responses. **A**–**C)** FC–FC score between model and data is plotted as a heat map in *G*–*B*^*e*^ space for the three model regimes considered here (see text): **A** ‘Fixed-point’ regime, **B** ‘Hysteresis’ regime, and **C** ‘Limit-cycle’ regime. Corresponding *J*^tot^–*E* bifurcation diagrams (cf. Eq. (4)) for each regime are shown in the right-hand panels **D**–**F**, showing stable *E* fixed points (solid), unstable *E* fixed points (dotted), and minima and maxima of limit-cycle oscillations (solid lines with shading). Dashed vertical lines represent the minimum *J*^tot^ corresponding to selected *B*^*e*^ values. Grey horizontal lines represent the range of *J*^tot^ values across regions and time for a sample simulation from the corresponding point in *G*–*B*^*e*^ space annotated in **A**–**C**. Parameter values for each regime are in Table S2.

We can understand the dynamics of an individual brain region in terms of the variation in its inputs over time, *J*_tot_(*t*) (recalling that *J*_tot_ is high when a region has many inputs from other high-activity, or high-*E*, regions). To understand this in more detail, we consider two key parameters that control the range of *J*_tot_ that can be explored by a given brain region. As per Eq. (4), these parameters are: (i) the excitatory firing threshold, *B*_*e*_, which contributes a constant offset to *J*_tot_; and (ii) the global coupling constant, *G*, which scales each region’s response to excitatory inputs from other connected regions. Increasing *B*_*e*_ decreases *J*_tot_ for all cortical regions, shifting the range of *J*_tot_ explored by network nodes to the left on the *J*_tot_–*E* bifurcation diagrams. We can see this from the annotated levels of input, *J*_tot_, corresponding to selected *B*_*e*_ values (when *J*_*e*_ = 0) as vertical dashed lines in Figs 2D–F. Low *G* ≈ 0 removes the effect of inter-regional coupling altogether (*J*_*e*_ ≈ 0), resulting in a very narrow range of *J*_tot_ around *B*_*e*_, while increasing *G* allows individual regions to respond more strongly to external inputs, and thus span a greater range of *J*_tot_ values. Given fixed values of *B*_*e*_ and *G*, the key factor controlling how different brain regions differ in *J*_tot_ in the homogeneous model is their connected neighbors, with high in-degree regions having more inputs and thus the potential to achieve a higher *J*_*e*_ and *J*_tot_ than low in-degree regions.

We are now able to analyze how different values of *B*_*e*_ (which adjusts the baseline of *J*_tot_) and *G* (which scales the excitatory inputs, *J*_*e*_, relative to this baseline) shape the dynamics of a given brain region. For example, some combinations of *B*_*e*_ and *G* confine all nodes in the network to a fixed-point attractor, whereas others allow some nodes to span one or multiple bifurcations. Different choices of *G* and *B*_*e*_ control the diversity of dynamical features supported by the model, but what types of configurations yield high FC–FC scores, *ρ*_FCFC_? Our results, comparing across a range of both *G* and *B*_*e*_, are shown as heat maps for the Fixed Point (Fig. 2A), Hysteresis (Fig. 2B), and Limit Cycle (Fig. 2C) regimes. To visualize the correspondence between points in *G*–*B*_*e*_ space and the resulting range of *J*_tot_(*t*) (and hence accessible dynamical regimes) they correspond to in the model simulation (range taken across time and nodes), we annotated this range in Figs 2D–F for key selected points in each corresponding heat map—labeled as ‘i’, ‘ii’, etc. For example, points towards the left of the *G*–*B*_*e*_ heat map correspond to low *G* and thus narrowing the range of *J*_tot_, while points near the top of the heat map correspond to high *B*_*e*_ and hence low baseline inputs; hence points labeled ‘i’ correspond to low and narrow ranges of *J*_tot_, as annotated to the bifurcation diagrams in Figs 2D–F.

We first note a wide range of *ρ*_FCFC_ in all cases, indicating that model performance depends strongly on the local response to inputs, *G* and *B*_*e*_, and thus the types of dynamical regimes available to the nodes of the coupled network. We also see that each dynamical regime exhibits characteristic regions of *G*–*B*_*e*_ space in which there is high FC–FC correspondence (colored red in Figs 2A–C), which reaches as high as *ρ*_FCFC_ = 0.52 (for the Fixed-Point regime), *ρ*_FCFC_ = 0.50 (Hysteresis regime), and *ρ*_FCFC_ = 0.56 (Limit-Cycle regime). All three model regimes can capture FC better than the direct correlation between SC and FC, *ρ*_SCFC_ = 0.42, indicating a benefit of accounting for distributed dynamics via coupled dynamical equations in capturing FC. Furthermore, model performance is consistent with, or higher than recently reported results for mouse cortex using a reduced Wong–Wang model (48) in a bistable regime (and using a Balloon–Windkessel BOLD filter (41) and a linear correlation, *ρ*_FCFC_): 0.35 ⪅ *ρ*_FCFC_ ⪅ 0.50 (24).

To understand how the model can produce high FC–FC, *ρ*_FCFC_ = 0.52 ± 0.03, in the Fixed Point regime (Fig. 2A), we start by exploring the qualitatively different types of input– output responses in Fig. 2D. At high excitatory firing threshold, *B*_*e*_, and low coupling, *G* (labeled ‘i’ in Figs 2A,D), nodes can only access the relatively flat, low-*E* steady-state branch, weakening inter-regional communication across the brain and leading to poor FC–FC. A similar suppression of inter-regional communication, and resulting low *ρ*_FCFC_, occurs when the model is confined to the upper branch at low *B*_*e*_ and high *G* (labeled ‘iii’ in Figs 2A,D). In the intermediate region, labeled ‘ii’ in Figs 2A,D, we obtain high FC–FC scores, up to a maximum *ρ*_FCFC_ = 0.52 ± 0.03 (at *B*_*e*_ = 3.3 mV, *G* = 0.65 mVs). Here, brain areas can access the sharp gradient of the sigmoid-like stable branch in *E*, and are thus highly sensitive in their response to variations in the activity of neighboring brain regions. This gives us the somewhat surprising result that this very simple model, in a regime in which regions respond to the aggregate activity of their neighbors (but without any complex local dynamical features like bifurcations or oscillations) can produce high *ρ*_FCFC_ = 0.52, consistent with results reported recently using more complex models (24). This is qualitatively consistent with direct structural connections providing a strong constraint on the resulting FC (33), with non-direct interactions providing a more minor perturbation (49).

We next investigated a ‘Saddle Node’ model regime (using parameters from (46)), that involves a pair of saddle-node bifurcations with an intermediate bistable region, shown in Figs 2B,E. We obtained qualitatively similar results to the Fixed-Point regime analyzed above: *ρ*_FCFC_ is low when nodes are confined to relatively flat low-*E* branch (at low *G* and high *B*_*e*_, labeled ‘i’ in Figs 2B,E) or the high-*E* branch (high *G* and low *B*_*e*_, labeled ‘iii’ in Figs 2B,E), where responses to external inputs are weak. Stronger FC–FC scores (e.g., a maximum *ρ*_FCFC_ = 0.50 ± 0.14 at *B*_*e*_ = 3.7 mV and *G* = 0.35 mVs) again arise in the intermediate region, where the local activity response is most sensitive to driving inputs, *J*_tot_ (labeled ‘ii’ in Figs 2B,E). The difference now is the increased diversity of supported dynamics: regions coexist between the stable low-*E* and high-*E* states, and can switch between them. This bistability leads to a greater dynamical repertoire of regions in the network, including longer-timescale switching (cf. Fig. S2), but this is not reflected in an improved *ρ*_FCFC_.

Finally, we investigated model dynamics in the neighborhood of a stable limit cycle, separated by two Hopf bifurcations (model parameters from Heitmann et al. (45)), shown as a *J*_tot_–*E* bifurcation diagram in Fig. 2F. As for the two regimes studied above, when nodes are confined to a relatively flat stable branch, labeled ‘i’ and ‘v’ in Figs 2C,F, FC– FC scores are low. For a similar reason, we also find low FC–FC when nodes are confined to a limit-cycle oscillation (labeled ‘iii’ in Figs 2C,F), where nodes have a restricted ability to respond to their inputs in a way that their neighbors can meaningfully respond to (since nodes are coupled via *E*, cf. Eq. (3)). But the heat map in Fig. 2C reveals two regions of *G*–*B*_*e*_ space with high *ρ*_FCFC_, labeled ‘ii’ and ‘iv’. In the region labeled ‘ii’ (e.g., *ρ*_FCFC_ = 0.38 ± 0.03 at *B*_*e*_ = 2.8 mV, *G* = 0.45 mVs), nodes sit on a stable branch which has a small but sufficient curvature to enable local activity to respond, albeit weakly, to inputs from connected regions. But the best fits to data, reaching *ρ*_FCFC_ = 0.56 ± 0.04 (at *B*_*e*_ = 1.5 mV, *G* = 0.7 mVs), are found in the region labeled ‘iv’ in Figs 2C,F. In this region of *B*_*e*_–*G* space, nodes can access two distinctive types of dynamics: the limit-cycle regime (at low *J*_tot_) and the high-*E* fixed-point attractor (at high *J*_tot_). High FC–FC scores are also obtained when nodes can also access the low-*E* stable branch (at high *G* and high *B*_*e*_). Importantly, the high-*E* branch at high *J*_tot_ has a relatively sharp dependence on *J*_tot_, a feature that is common to obtaining high-*ρ*_FCFC_ scores in all three model regimes. Together, our analyses in this section demonstrate the importance of a model that allows nodes to respond sensitively to inputs from their network neighbors for reproducing FC.

#### Interpreting simulated dynamics in terms of bifurcation diagrams

Bifurcation diagrams provide an understanding of the dynamical regimes accessed by individual nodes, and the way in which they respond to changes in inputs, information that can guide understanding of the complex distributed dynamics that result from a full model simulation. For the Limit-Cycle regime, simulated multivariate time series and corresponding FC matrices are plotted in Fig. 3 for points labeled ‘ii’, ‘iii’, and ‘iv’ in Figs 2C,F. In ‘ii’, nodes are confined to the low-*E* stable branch and, accordingly, the dynamics consist of noisy deviations from a low-*E* stable fixed point (Fig. 3D). These perturbations can drive changes in structurally connected nodes, yielding weak pairwise correlations shown in Fig. 3A. In ‘iii’, when nodes are mostly confined to the limit cycle and *ρ*_FCFC_ is low, most nodes exhibit oscillations (with some longer-timescale deviations, cf. Fig. 3E), and a minority of other nodes (situated near the low-*J*_tot_ Hopf bifurcation) move between noisy deviations from the low-*E* stable branch and oscillatory limit-cycle dynamics. This results in very high pairwise correlations between groups of synchronized oscillatory nodes, *r >* 0.8, such that the underlying structural connections play less of a role in shaping the pairwise correlation structure, resulting in a low FC–FC score. In ‘iv’, with the highest *ρ*_FCFC_, the Hopf bifurcations facilitate complex spatiotemporal dynamics shown in Fig. 3F. While many nodes spend most of the simulation near the high-*E* stable branch (those with high *J*_tot_), we observe periods of time during which groups of nodes (near the Hopf bifurcation) display synchronized oscillations, embedded in globally complex and distributed dynamics on longer timescales. These analyses demonstrate how analyzing the response of local nodes to inputs, as ranges of *J*_tot_ in a bifurcation diagram (as in Figs 2D–F), can ground an understanding of the complex distributed dynamics that result from the full coupled model, which can be visualized effectively as heat maps (Fig. 3).

**Fig. 3.**
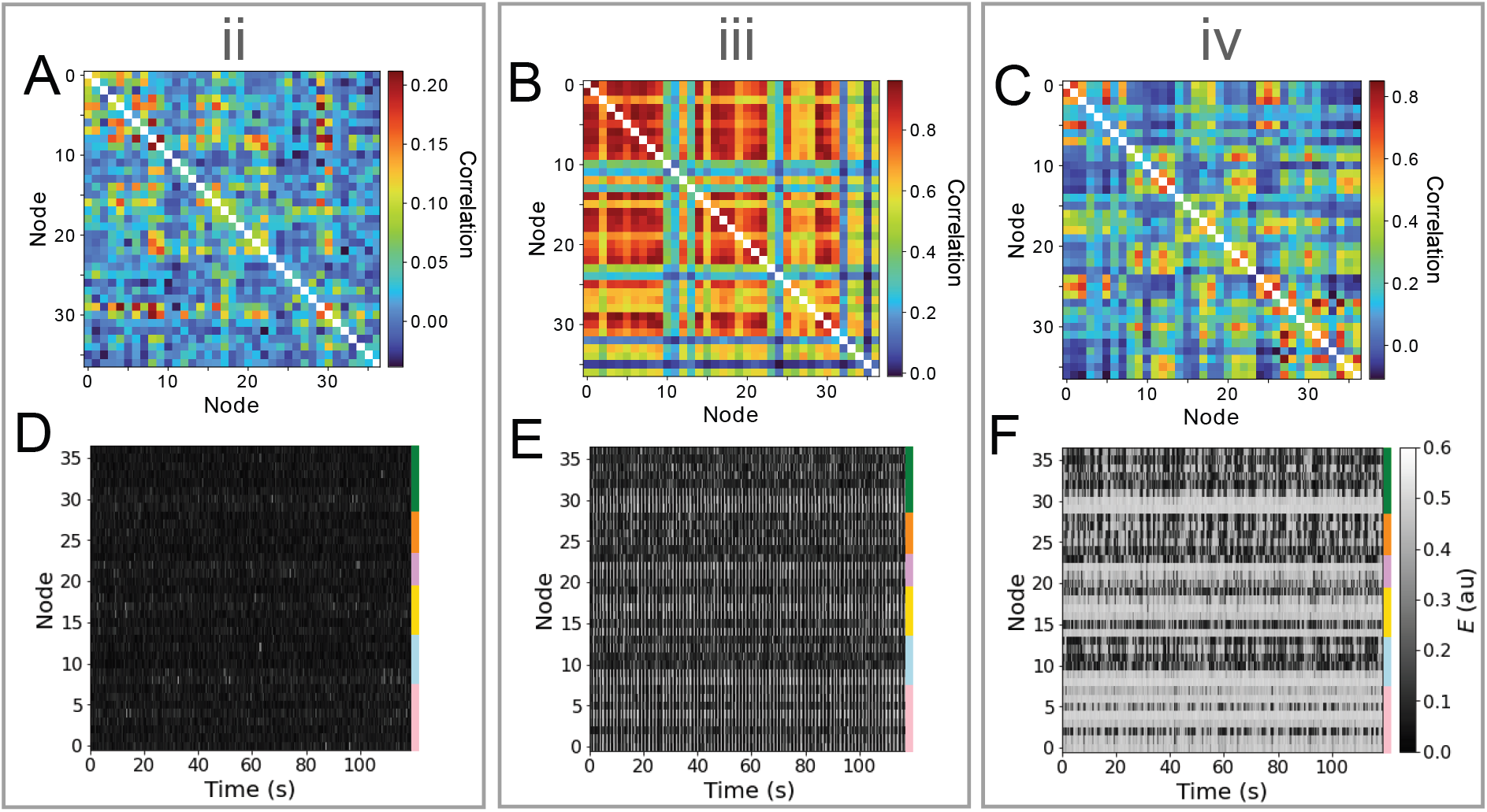
Different dynamical features of the limit-cycle regime yield very different dynamics, including noisy deviations about a stable fixed point, synchronous oscillations, and a complex distributed dynamics featuring intermittent synchronization with high FC–FC. Here we investigate simulated time series (lower row) and functional connectivity matrices (upper row) for three regions in *B*^*e*^–*G* space annotated ‘ii’, ‘iii’, and ‘iv’ in Fig. 2. **A**–**C** Simulated functional connectivity matrices are plotted for ‘ii’, ‘iii’, and ‘iv’, respectively. **D**–**F** Simulated *E* time series are plotted as a node *×* time heat map (or ‘carpet plot’ (50)) for all brain regions for ‘ii’, ‘iii’, and ‘iv’, respectively. Colored bars label the six cortical divisions listed in Table S1. In all plots, nodes are ordered as per Table S1.

#### Resolving inter-regional differences in inputs

The variation in qualitative dynamics across individual brain areas in the multivariate time series plotted in Figs 3D–F indicates that different network elements are accessing different dynamical regimes permitted by the model, resulting from substantial variability in the *J*_tot_(*t*) experienced by different nodes. Since all nodes are governed by the same dynamical rules, and hence the same bifurcation diagrams, we can annotate *J*_tot_(*t*) ranges onto a common bifurcation diagram to understand how the dynamics of individual regions are governed by different types of inputs from their connected neighbors. That is, rather than plotting just the overall range of *J*_tot_ (from the minimum to the maximum across all nodes), as in Figs 2D– F, we can resolve the individual ranges of *J*_tot_ experienced by each individual node on the *J*_tot_–*E* bifurcation diagram. An example is shown for the Limit-Cycle regime at ‘iv’ in Fig. 4A (where we have plotted *J*_*e*_ instead of *J*_tot_, equivalently, for a fixed *B*_*e*_ = 1.5 mV, cf. Eq. (4)). We see how, even with fixed dynamical rules, the range of *J*_*e*_ experienced by individual nodes varies markedly. Some regions have low *J*_*e*_ across the simulation, like the dorsal retrosplenial area, RSPd (annotated in Fig. 4A), and therefore only display oscillations, as plotted in Fig. 4B. Other regions with high *J*_*e*_ across the simulation, like the posterior parietal association areas, PTLp (annotated in Fig. 4A), are confined to the stable high-*E* branch across the full simulation and display dynamics consistent with noisy deviations from a fixed point, as shown in Fig. 4B. Regions like the ventral retrosplenial area, RSPv (annotated in Fig. 4A), span the Hopf bifurcation, and thus exhibit more complex patterns that contain both oscillatory dynamics and noisy excursions about a stable fixedpoint, depending on fluctuations in inputs, *J*_*e*_(*t*). The short samples of *E*(*t*) for six annotated Medial regions in Fig. 4B reveal some of these dynamics, including dynamic phase relationships between the oscillatory populations. These findings demonstrate the usefulness of interpreting the dynamics of coupled mass models in terms of time-varying inputs to the constituent populations.

**Fig. 4.**
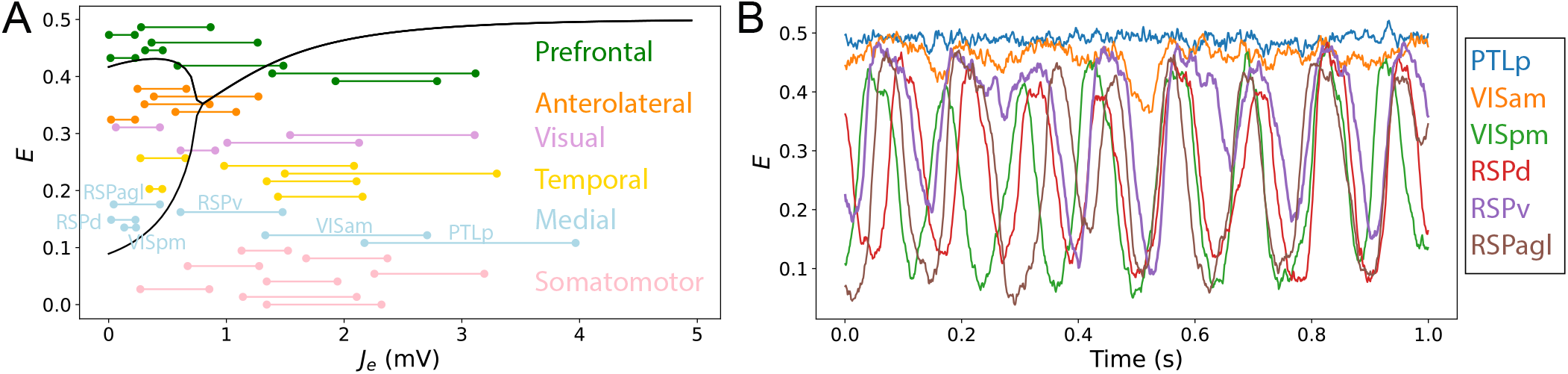
Resolving different ranges of inputs, *J*^*e*^, **experienced by different network nodes allows us to understand their variable dynamical behavior in a coupled network model**. Here we focus on the point labeled ‘iv’ in the Limit-Cycle regime (Fig. 2C,F), *B*^*e*^ = 1.5 mV, *G* = 0.7 mVs, in which nodes differ substantially in their inputs, *J*^*e*^, and hence their resulting dynamics. **A** Bifurcation diagram for *E* and a function of *J*^*e*^ (as Fig. 2F), with ranges of net excitatory drive, *J*^*e*^, across the model simulation annotated for each brain region (colored according to the six labeled divisions). All regions are ordered according to Table S1, and are labeled for the six Medial regions, which are plotted in **B. B** *E* time series for the six Medial regions—PTLp, VISam, VISpm, RSPd, RSPv, and RSPagl—shown for the final 1 s of the simulation.

### Understanding heterogeneity in local dynamical rules

Above, we used bifurcation diagrams to show that complex distributed dynamics in a neural-mass model can be understood in terms of the responses of individual regions to inputs from their connected neighbors. Despite equivalent local dynamical rules, and hence identical bifurcation diagrams for all brain regions, we found substantial inter-regional variability in accessible dynamical regimes and resulting activity dynamics, due to differences in structural connectivity and resulting *J*_*e*_(*t*). In this section, we aim to understand the effect of varying the local dynamical rules themselves, by incorporating spatial heterogeneity in the properties of local cortical circuits (via a corresponding variation in model parameters). Specifically, we varied excitatory and inhibitory coupling strengths of individual brain areas according to excitatory and inhibitory cell-density data (4). We focused on the Limit Cycle regime of the W–C model described above, which displayed the richest dynamical repertoire and highest *ρ*_FCFC_. As described in Methods, we used relative variations in excitatory and inhibitory cell densities across cortical areas to define a corresponding variation in *R*_*e*_ and *R*_*i*_, which proportionally adjust coupling parameters—*w*_*ii*_, *w*_*ie*_, *w*_*ei*_, *w*_*ee*_—across brain areas. Setting *R*_*e*_ = *R*_*i*_ = 0 for all areas recovers the homogeneous model studied above (see Eq. (5) for details). This simple formulation allows us to understand how varying the excitatory and inhibitory coupling parameters across areas, in accordance with underlying excitatory and inhibitory cell densities, shape the dynamical responses of individual areas to inputs, and hence the resulting model dynamics.

#### Levels of excitation and inhibition perturb bifurcation diagrams

To understand how variations in *R*_*e*_ and *R*_*i*_ affect model dynamics, we first analyze how these parameters shape the *J*_tot_–*E* bifurcation diagrams for an individual area. The effect of *±*10% variations to coupling parameters (corresponding to the ranges *™*0.1 *< R*_*e*_ *<* 0.1 and *™*0.1 *< R*_*i*_ *<* 0.1), are shown as *J*_tot_–*E* bifurcation diagrams in Fig. 5, varying just *R*_*e*_ (Fig. 5A), just *R*_*i*_ (Fig. 5B), *R*_*e*_ and *R*_*i*_ together with *R*_*e*_ =™*R*_*i*_ (Fig. 5C), and *R*_*e*_ and *R*_*i*_ such that *R*_*e*_ = − *R*_*i*_ (Fig. 5D). We find that even these relatively small, ≈ 10%, perturbations have a substantial effect on the dynamical responses of individual areas, affecting: (i) the range of *J*_tot_ over which model exhibits stable oscillations; (ii) the oscillation amplitudes themselves; and (iii) steady-state activity levels. As shown in Fig. 5, cortical areas with a higher excitatory cell density, *R*_*e*_, have higher-amplitude oscillations, a wider range of *J*_tot_ over which stable oscillations are exhibited, and, for the same *J*_tot_, increased activity, *E*, in the upper branch. Different changes result from modifying the inhibitory cell density, shown in Fig. 5B: increasing *R*_*i*_ shifts the same bifurcation and fixed-point structure to higher *J*_tot_ (equivalent to raising the firing threshold, *B*_*e*_). That is, regions with higher inhibitory cell density, *R*_*i*_, require a greater aggregate input, *J*_*e*_, to produce the same dynamics. Varying both *R*_*e*_ and *R*_*i*_, shown in Figs 5C,D, yields combinations of the individual perturbations from *R*_*e*_ and *R*_*i*_ individually. These results demonstrate how relatively small variations in excitatory and inhibitory coupling parameters can have large effects on the bifurcation structure and dynamical regimes exhibited by local cortical regions. The effects are more dramatic for *R*_*e*_ and *R*_*i*_ values in the range from 0.5 to 0.5, where the Hopf bifurcations can be removed altogether from the Limit Cycle regime (Fig. S3), or additional stable states can be added via saddle-node bifurcations in the Hysteresis regime (Fig. S4).

**Fig. 5.**
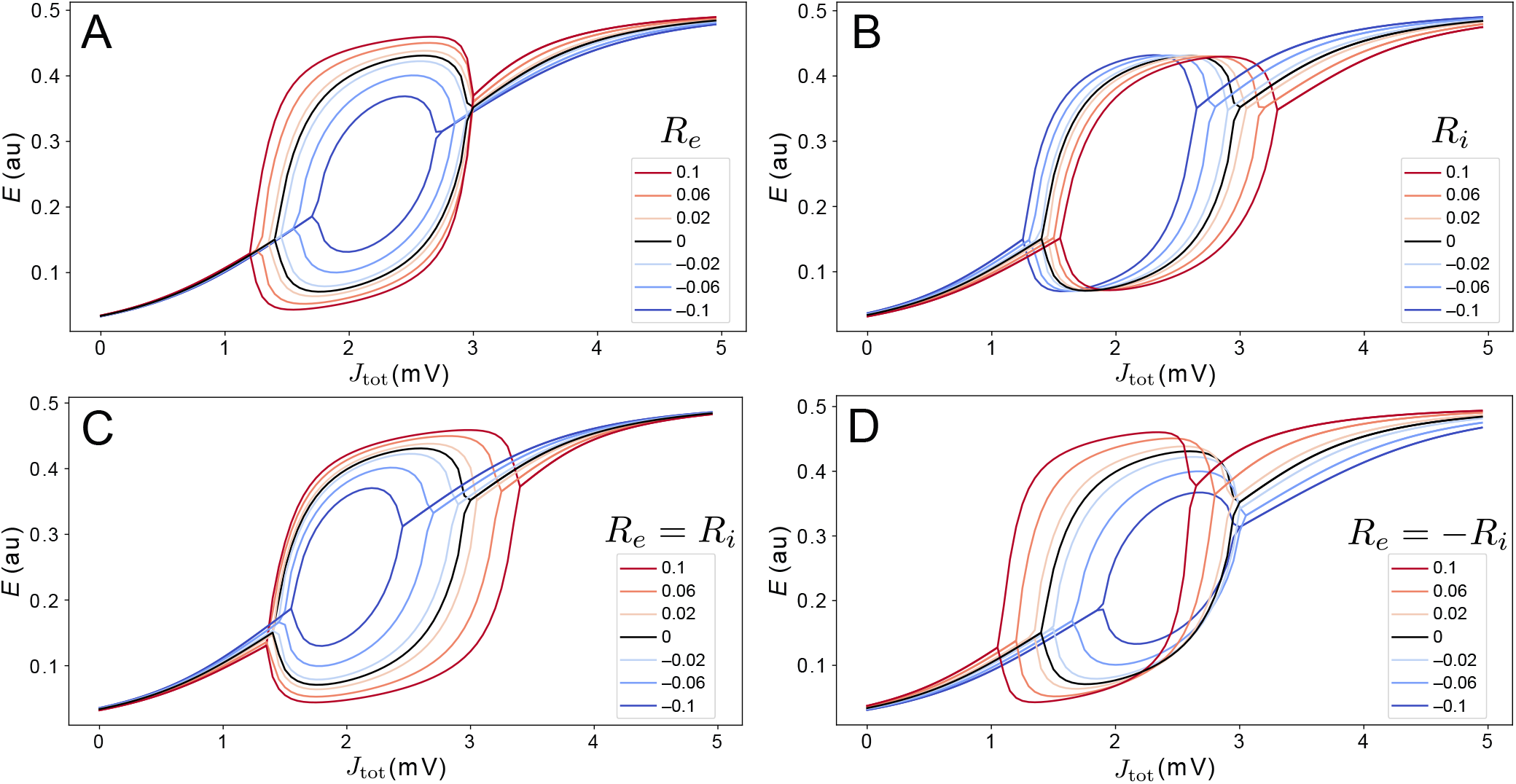
Variations in excitatory and inhibitory cell density modify the dynamical regimes accessible to cortical regions. We model the effect of variations in excitatory and inhibitory cell density via perturbation parameters *R*^*e*^ and *R*^*i*^, respectively, as defined in Eq. (5). Relative to the nominal bifurcation diagram, *R*^*e*^ = *R*^*i*^ = 0 (black), we investigate variations in *™*0.1 *≤ R*^*e*^ *≤* 0.1 and *™*0.1 *≤ R*^*i*^ *≤* 0.1. Four types of variation were investigated: **A** *R*^*e*^ only (*R*^*i*^ = 0); **B** *R*^*i*^ only (*R*^*e*^ = 0); **C** *R*^*e*^ and *R*^*i*^, such that *R*^*e*^ = *R*^*i*^; and **D** *R*^*e*^ and *R*^*i*^, such that *R*^*e*^ = *™R*^*i*^. The legend indicates values of *R*^*e*^.

#### Understanding mouse cortical model dynamics constrained by excitatory and inhibitory cell densities

In the heterogeneous model, individual different brain areas differ both in their *J*_tot_ values that they receive from their coupled neighbors (due to differences in their structural connections), but also have different dynamical rules, due to different individual combinations of *R*_*e*_ and *R*_*i*_ values. As demonstrated above for the homogeneous model, this understanding of the dynamical responses of individual brain areas to inputs from across the network is crucial to guiding understanding of the complex, distributed dynamics of the full coupled model. In this section, we explore how the impact of local variations in *R*_*e*_ and *R*_*i*_ can be visualized and used to understand the dynamics of the full coupled model. Recall that our heterogeneous model is formulated with a single new parameter, *σ*, that defines how strongly relative differences in excitatory and inhibitory cell densities are mapped to corresponding changes in the model’s coupling parameters (Eq. (6)). For the Limit-Cycle regime, we investigated how FC–FC scores change as we introduce a greater degree of inter-areal heterogeneity, *σ*. The variation in *ρ*_FCFC_ as a function of *σ* across the range 0 ≤ *σ* ≤ 1 is shown in Fig. 6E. We did not find a substantial increase in *ρ*_FCFC_ when incorporating heterogeneity, *σ >* 0, although there was a modest improvement relative to the homogeneous model (*σ* = 0) for *σ* = 0.2, yielding *ρ*_FCFC_ = 0.60 0.05. Testing this result against a null distribution (obtained by repeating the procedure but with randomly permuted excitatory and inhibitory cell-density data) using a permutation test yielded *p* ≈ 0.15, indicating that *ρ*_FCFC_ = 0.60 does not constitute a significant improvement relative to the homogeneous model (see Sec. S2 for details). As we discuss later, this result may be contributed to to the small number of regions in the model, the simplicity of the dynamical equations, or the dominance of SC in constraining FC in the anesthetized mouse (33).

**Fig. 6.**
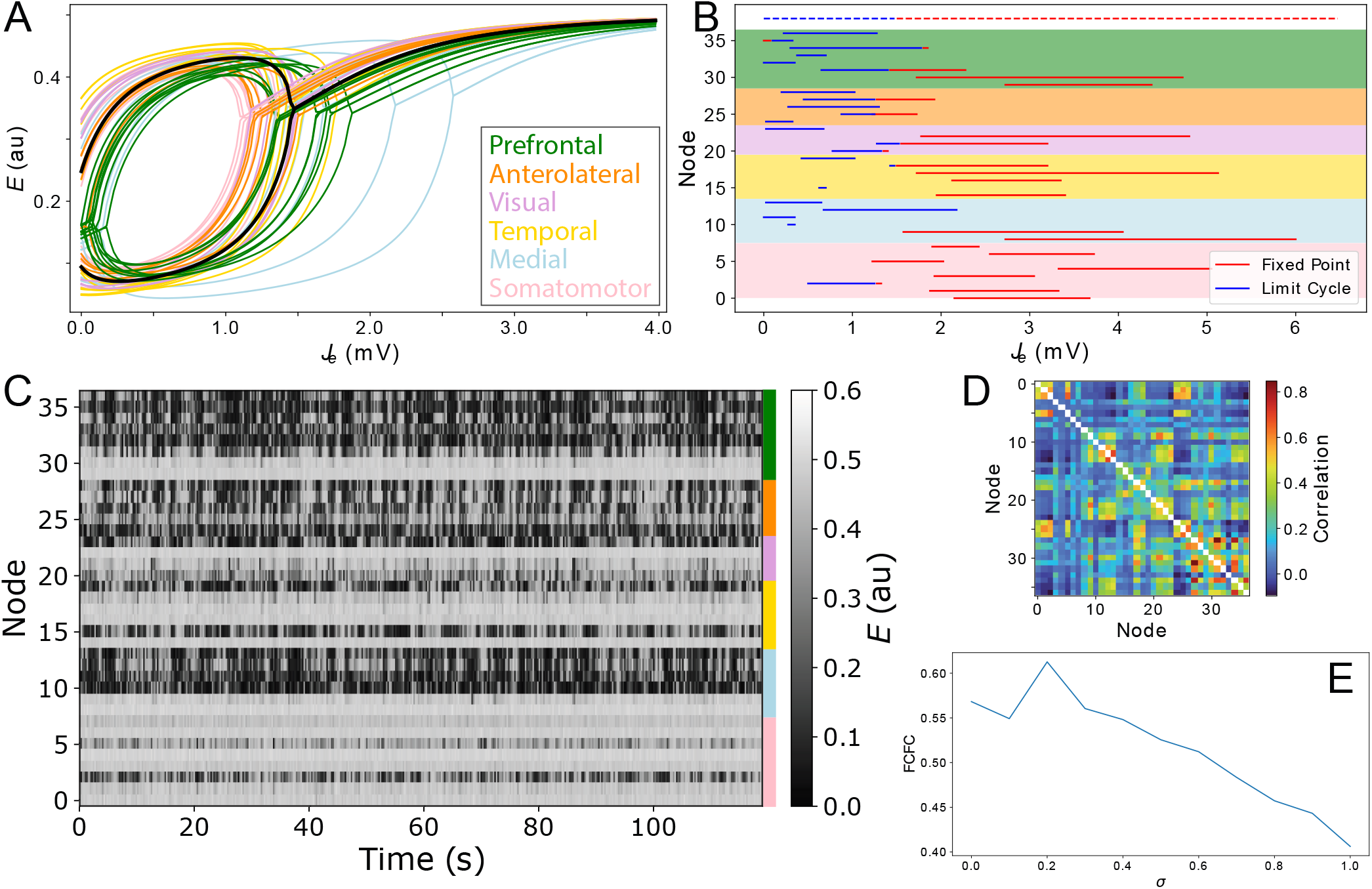
Modeling spatial variation in local excitatory and inhibitory cell densities produces complex distributed dynamics. **A** Bifurcation diagrams are plotted for all cortical areas according to their excitatory and inhibitory cell densities. Regions are colored according to their labeled anatomical grouping and the homogeneous case (*R*_*e*_ = *R*_*i*_ = 0) is shown in black for comparison. **B** The type of equilibrium dynamics displayed by a given cortical region, limit cycle (blue) or fixed point (red), is plotted as a function of *J*_*e ™*_ *B*_*e*_ for all cortical regions for the range of *J*_*e ™*_ *B*_*e*_ they experience across the model simulation. Nodes are ordered as per Table S1 and shading reflects the six anatomical groupings labeled in A. Dashed lines shown at the top correspond to the uniform case (*R*_*e*_ = *R*_*i*_ = 0) for comparison. **C** Simulated time series for all brain areas are plotted as a heat map. Colors annotated to the right label the six anatomical groupings listed in A. **D** Simulated functional connectivity matrix. **E** FC–FC score as a function of the scaling parameter, *σ*, Eq. (6).

While we did not find evidence of a significant improvement in FC–FC score from a simple incorporation of heterogeneity, our main aim was to demonstrate how tools from dynamical systems can help to understand the complex coupled dynamics of such a spatially heterogenous model. We used the point, *σ* = 0.2, inferred above as a suitable example for this purpose. With *σ* = 0.2, we plotted *J*_*e*_–*E* bifurcation diagrams for all brain regions on the same plot in Fig. 6A. The plot shows how differences in excitatory and inhibitory cell densities results in different bifurcation diagrams, that correspond to similar qualitative changes as analyzed in Fig. 5 above. Specifically, brain regions now differ substantially in their: critical values, *J*_*e*_, that separate limit cycle from fixedpoint dynamics; ranges of *J*_*e*_ in which oscillations are stable; oscillation amplitudes; and fixed-point activity levels, *E*, in the upper branch (for a given *J*_*e*_). Compared to the homogeneous model, two regions with the same input, *J*_*e*_, no longer indicates that they will be subject to the same dynamical rules.

To understand how these changes in local dynamical rules affect the resulting cortical dynamics, we next plotted the range of *J*_*e*_ that each node experiences across the simulation. As shown in Fig. 6B, this can be represented as a horizontal line, distinguishing *J*_*e*_ values corresponding to what stable dynamical feature—’limit cycle’ or ‘fixed point’—according to each region’s individual bifurcation structure. This results in a richer dynamical landscape for the model: some brain regions can access both stable limit cycle and fixed-point dynamics, others can only access the high-*E* fixed-point equilibrium, while others can access just the limit-cycle attractor. It is useful to connect the range of dynamical regimes each region accesses across the simulation, shown in Fig. 6B, with the *E* dynamics themselves, shown in Fig. 6C. We can clearly see the high-*E* regions on the upper stable branch, as well as the more complex intermittent oscillations of regions that can access limit-cycle dynamics. The functional connectivity matrix from this simulation is shown in Fig. 6D. This representation of pairwise correlations in the model dynamics hides much of the richness of the individual time series themselves (Fig. 6C), and the dynamical rules that underlie them (Figs 6A,B). The ability to represent qualitative dynamical regimes of individual regions in a coupled network model— as *J*_*e*_–*E* bifurcation diagrams with individual ranges of *J*_*e*_ explored for each region—provides a powerful illustration of the dynamics supported by the coupled components of a complex networked dynamical model.

## Discussion

In this paper we developed a neural-mass model of the mouse cortex. We showed how bifurcation diagrams can be used to understand how regional differences in dynamics result from differences in inputs, *J*_tot_, and delineated the types of dynamical regimes that yield good fits to experimental functional connectivity. We first analyzed a homogeneous model in which all regions are governed by identical dynamical rules to show how regional variations in dynamics result from differences in inputs (driven by differences in structural connectivity). We then extended this treatment to a heterogeneous model in which the bifurcation structures themselves vary across regions due to variation in local excitatory and inhibitory cell densities. Our results provide a useful framework for understanding the mechanisms that underlie complex simulated model dynamics, using a combination of local bifurcation diagrams (annotated with ranges of inputs for different regions) and visualizations of the multivariate timeseries dynamics (as ‘carpet plots’ (50)). These analyses will be particularly important for understanding how the brain’s microscale circuits give rise to the complex distributed dynamics observed in brain-imaging experiments.

A common scientific goal of modeling a system is to accurately reproduce important properties of it, while also gaining an understanding of how it does so. While successful approaches have been demonstrated for maximizing goodness of fit (sometimes optimizing large numbers of parameters (18, 19, 51)), obtaining understanding is a key challenge for complex nonlinear models of brain dynamics. The analyses and visualizations demonstrated in this work aim to provide an understanding of the model dynamics in terms of the dynamical regimes that individual regions can access, shaped by their inputs from coupled neighbors. Key analyses include: (i) assessing the role of input parameters *B*_*e*_ and *G* in shaping empirical FC fits in terms of corresponding ranges spanned across *J*_*e*_–*E* bifurcation diagrams (Fig. 2); (ii) annotating of *J*_*e*_ for all cortical regions onto a common bifurcation diagram (Fig. 4A); (iii) analyzing perturbations to bifurcation diagrams due to variations in local circuit properties (Fig. 5); and (iv) annotating region-specific qualitative equilibrium dynamics across ranges of *J*_*e*_ for all regions in a single plot (Fig. 6B). As whole-brain models develop to incorporate whole-brain datasets—including whole-brain maps of gene-expression and cell types (2, 52)—these types of analyses will be crucial to understanding how this complexity shapes the underlying dynamical mechanisms, both at the level of individual brain regions, and their distributed interactions.

While many studies focus on determining an optimal working point, i.e., structural connectome scaling *G*, we find that the offset (*B*_*e*_ in the present model), is also critical to determining how local regions respond to inputs, and hence the resulting *ρ*_FCFC_. We also found strong fits to empirical FC whenever brain regions were able to respond to inputs with sufficient gain, likely reflecting the strong role of direct structural connections in shaping FC in the anesthetized mouse (33). In particular, even in the Fixed-Point regime, in which the model exhibits the most constrained dynamical repertoire, we report *ρ ≈* 0.52, consistent with other published results in the literature (FC–FC scores up to ≈ 0.5 (24) using a Wong–Wang model (48)). Only a small improvement was found when the model operated near a Hopf bifurcation, *ρ*_FCFC_ = 0.56. This highlights the ability of simple dynamical features to capture aspects of measured dynamics, consistent with prior comparisons demonstrating high performance of simple models (53, 54). The results also demonstrate the importance of comparing model performance against simpler benchmarks, and justifying increased model complexity only if it accompanies enhanced explanatory power.

Incorporating spatial variations in local dynamical rules according to whole-brain maps has immense potential in allowing us to connect new physiological understanding of neural circuits to the whole-brain dynamics that they enable. In this work, we incorporated spatial variations in excitatory and inhibitory cell densities as a corresponding perturbation to coupling parameters between *E* and *I* populations, with a single scaling parameter, *σ*. However, there are alternative ways in which this heterogeneity could be implemented and constrained in future, for example, by allowing *σ* to differ for excitatory (*σ*_*e*_) and inhibitory (*σ*_*i*_) populations. Incorporating more detailed physiological data into correspondingly more complex biophysical models (e.g., incorporating multiple inhibitory cell types), brings further parametric freedom that needs to be properly constrained from a combination of physiological and neuroimaging data. Our approach for assessing the improvement in *ρ*_FCFC_ after incorporating celldensity data involved a permutation approach against randomized assignment of the data to regions (preserving the match between *e* and *i*, but permuting their assignment to brain regions), and did not reveal a significant improvement relative to null gradients (*p ≈* 0.15). This may be due to the relatively small number of brain regions included, and the focus on FC–FC as an evaluation metric rather than a more comprehensive set of evaluations. Other ways of assessing the improvement of the spatially heterogeneous model could also be explored, such as testing the *e* : *i* ratio against alternative spatial gradients (as Deco et al. (17)), or taking an optimization approach to estimate the optimal *e* and *i* gradients, and then assess their similarity to the measured excitatory and inhibitory cell-density data (as Wang et al. (19)).

There are many limitations of our general modeling approach that can also be extended in future work. First, we have focused here on a specific simple biophysical model, the Wilson–Cowan model (34, 35, 38), that allowed us to incorporate variations in excitatory and inhibitory cell-density data. We have focused on the behavior of the model in three specific dynamical regimes (a fixed point with gain, hysteresis, and limit cycle), but the results should be qualitatively applicable to those same dynamical regimes of other models. However, we note that other models with different dynamical features may display different behavior, such as the Wong–Wang model (16, 17, 19, 21, 48, 55), or models that incorporate cortico–thalamic interactions (25, 35, 56, 57). We also note that while our aim here was to understand the model dynamics directly, it is common practice to simulate a hemodynamic response, such as the Balloon–Windkessel model (41) or a more sophisticated hemodynamic response function (58). Simulating a slower hemodynamic response would introduce challenges in mapping bifurcation diagrams in *E* to the corresponding BOLD dynamics, and could lead to substantial differences to dynamics and resulting FC (e.g., in a bistable switching regime). As a result, our specific conclusions about model performance in different dynamical regimes may not generalize to different choices of hemodynamic responses. We also highlight our relatively simple treatment of structural connectivity, as a binary adjacency matrix, even though estimates of axonal connectivity strengths vary over at least four orders of magnitude (6). However, it remains an open question what greater structural connection ‘strengths’ (approximated by the number of axons connecting two brain areas), corresponds to dynamically, e.g., faster connection speeds, a stronger effect on local population mean dynamics, or some alternative type of response. We next note a major simplifying assumption in using a neural-mass model, which involves representing the spatially continuous cortical sheet as a set of 37 discrete cortical areas, abstracted away from their physical embedding (59). Given the spatial resolution of modern mouse-brain maps, and the often continuous spatial variation they reveal, it will be important to develop models that accurately capture this physical continuity, e.g., using a neural field approach (60).

Finally, we highlight the limitation of evaluating our model with respect to its ability to match only the linear correlation structure, FC, of the empirical dataset. fMRI data have a much richer dynamical structure than is captured by the static FC, including the dynamics of FC across a recording (16, 17, 43, 44) and the organization of regional timescales (8, 9). For example, despite producing very different patterns in simulated time series, we found similar fits, *ρ*_FCFC_, across the Fixed-Point, Hysteresis, and Limit-Cycle regimes of our homogeneous model, and when incorporating heterogeneity. The more complex distributed dynamics, including intermittent synchronization seen in carpet plots from the Limit Cycle regime (Fig. 3F) and when incorporating regional heterogeneity (Fig. 6C), qualitatively match the types of patterns seen in empirical fMRI dynamics better than in the fixed-point regime. This highlights the simplicity of the FC–FC score, *ρ*_FCFC_, in capturing only the pairwise linear correlation structure in the data, and indicates the need for future work to perform a more comprehensive evaluation. This should include similar visualizations of model performance across *B*_*e*_–*G* space (as Figs 2A–C), where the most distinctive models features for reproducing a greater range of characteristics of fMRI dynamics may be more clearly distinguished.

With the increasing availability of high-resolution neuroscience data, in space and time, the need for tools to provide interpretable accounts of their dynamics is pressing. Our work demonstrates a range of useful tools to analyze the behavior of coupled dynamical models of brain dynamics, helping them to provide understanding of the dynamical mechanisms that underpin their predictions. Our results emphasize the importance of benchmark comparison (e.g., a simple fixed-point model yields high FC–FC), visualization (e.g., very different dynamical patterns exhibited in carpet plots can yield similar correlation structures in FC), and proper statistical testing (e.g., while the heterogeneous model yields improved FC–FC, it is not significantly better than repeating the process on randomized data), practices that may help guide progress in the field.

