## Supplementary text, tables, and figures for "Extracting dynamical understanding from neural-mass models of mouse cortex"

### S1 Data

**Structural connectivity** The structural connectivity matrix used here was taken from the Allen Mouse Brain Connectivity Atlas [1]. We first reduced the original $213 \times 213$ connectivity matrix to a $37 \times 37$ matrix including our set of cortical areas. We then transformed it into an unweighted, directed connectivity matrix by retained only edges with a $p$-value less than 0.05 from the regression model fitted by Oh et al. [1].

**Cell-density data** We used excitatory and inhibitory cell-density estimates from Erö et al. [2]. To generate these data, the authors algorithmically generated cell positions and cell types for the entire mouse brain using transcriptional markers from the Allen Mouse Brain Atlas [3].

**Resting-state fMRI** fMRI data for 100 wild-type mice are taken from Zerbi et al. [4]. fMRI data consisted of blood-oxygen-level–dependent (BOLD) signals recorded from 100 anesthetized mice measured at rest for a period of 15 min using a Biospec 70/16 small animal MR system operating at 7T, equipped with a cryogenic quadrature surface coil for signal detection (Bruker BioSpin AG, Fällanden, Switzerland). The data were parcellated using the Allen Common Coordinate Framework (CCF v3). We took time-series data from the 37 cortical regions analyzed here, and computed a functional connectivity matrix for each mouse as pairwise Pearson correlations. These matrices were averaged across mice to yield a group-average FC that was used as the basis of comparison for computing FC–FC scores.

### S2 Permutation Testing

We aimed to perform a simple statistical test to assess the improvement in FC–FC score resulting from incorporating spatial heterogeneity, via excitatory and inhibitory cell densities, into a coupled network model of W–C neural masses. In particular, our method involves testing for an improved $\rho_{\mathrm{FCFC}}$ multiple times, across a range of $\sigma$, and therefore has a greater potential to find an improved FCFC, even in the absence of a robust underlying signal. We assessed the statistical significance of the measured result, $\rho_{\mathrm{FCFC}} = 0.60$ (at $\sigma = 0.2$), relative to a null model in which excitatory and inhibitory cell densities were assigned to regions at random. This was done by randomly permuting the rows of the $38 \times 2$ (region $\times$ cell density) matrix, and thus does not destroy excitatory—inhibitory correlation structure. We estimated a $p$-value for the result $\rho_{\mathrm{FCFC}} = 0.60$ as a permutation test relative to a null distribution from 100 randomized simulations, returning the maximum FC–FC score across the range $0 \leq \sigma \leq 1$ in each case. This procedure yielded the estimate $p \approx 0.15$.

| Functional Group | Number | Acronym | Region Name |
|---|---|---|---|
| Somatomotor | 0 | SSs | Supplemental somatosensory area |
| (Pink) | 1 | MOp | Primary motor area |
| | 2 | SSp-n | Primary somatosensory area, nose |
| | 3 | SSp-ll | Primary somatosensory area, lower limb |
| | 4 | SSp-bfd | Primary somatosensory area, barrel field |
| | 5 | SSp-m | Primary somatosensory area, mouth |
| | 6 | SSp-tr | Primary somatosensory area, trunk |
| | 7 | SSp-ul | Primary somatosensory area, upper limb |
| Medial | 8 | PTLp | Posterior parietal association areas |
| (Light Blue) | 9 | VISam | Anteromedial visual area |
| | 10 | VISpm | posteromedial visual area |
| | 11 | RSPd | Retrosplenial area, dorsal part |
| | 12 | RSPv | Retrosplenial area, ventral part |
| | 13 | RSPagl | Retrosplenial area, lateral agranular part |
| Temporal | 14 | AUDd | Dorsal auditory area |
| (Gold) | 15 | AUDp | Primary auditory area |
| | 16 | AUDv | Ventral auditory area |
| | 17 | PERI | Perirhinal area |
| | 18 | TEa | Temporal association areas |
| | 19 | ECT | Ectorhinal area |
| Visual | 20 | VISal | Anterolateral visual area |
| (Plum) | 21 | VISp | Primary visual area |
| | 22 | VISl | Lateral visual area |
| | 23 | VISpl | Posterolateral visual area |
| Anterolateral | 24 | VISC | Visceral area |
| (Dark Orange) | 25 | GU | Gustatory areas |
| | 26 | AId | Agranular insular area, dorsal part |
| | 27 | AIv | Agranular insular area, ventral part |
| | 28 | AIp | Agranular insular area, posterior part |
| Prefrontal | 29 | MOs | Secondary motor area |
| (Green) | 30 | ACAd | Anterior cingulate area, dorsal part |
| | 31 | ORBl | Orbital area, lateral part |
| | 32 | PL | Prelimbic area |
| | 33 | ORBvl | Orbital area, ventrolateral part |
| | 34 | ORBm | Orbital area, medial part |
| | 35 | ACAv | Anterior cingulate area, ventral part |
| | 36 | ILA | Infralimbic area |

Table S1: **The 37 cortical regions modeled here.** Regions are listed by their ordering used in many plots in the main text, and grouped into six anatomical divisions from Harris et al. [5], with colors used for annotation in main text figures.

| | Regime | | |
|---|---|---|---|
| Parameter (units) | Fixed Point | Hysteresis | Limit Cycle |
| $w_{ee}$ (V s) | 12 | 16 | 11 |
| $w_{ei}$ (V s) | 15 | 12 | 10 |
| $w_{ie}$ (V s) | 10 | 10 | 10 |
| $w_{ii}$ (V s) | 8 | 3 | 1 |
| $b_i$ (mV) | 4 | 3.7 | 2.8 |
| $\tau_e$ (ms) | 10 | 10 | 10 |
| $\tau_i$ (ms) | 10 | 10 | 65 |
| $a_e$ ($V^{-1}$) | 1 | 1.3 | 1 |
| $a_i$ ($V^{-1}$) | 1 | 2 | 1 |

Table S2: **Parameter values corresponding to the three key model regimes studied in this work.** The 'Fixed Point' regime uses parameters modified from Sanz-Leon et al. [6] (modified to obtain a fixed-point), 'Hysteresis' regime uses parameters from Borisyuk and Kirillov [7], and the 'Limit Cycle' regime uses parameters from Heitmann et al. [8].

### S3 Figures

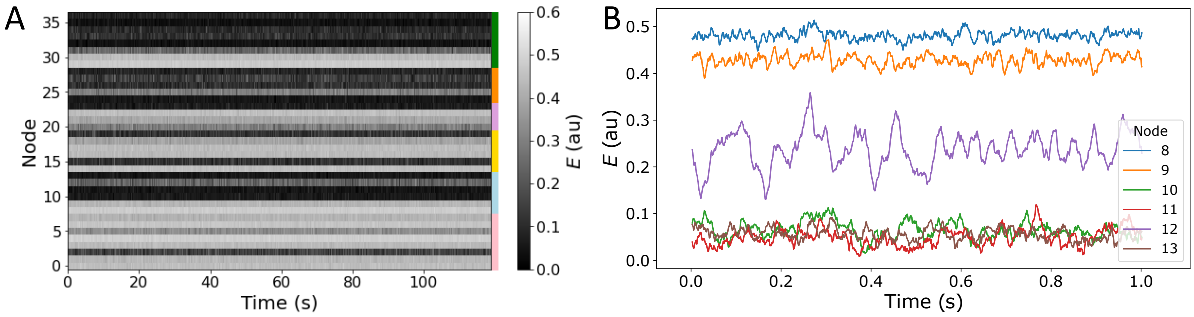


Figure S1: **Fixed-point simulation**. For the Fixed-Point regime model with the best FC–FC fit ($G = 0.65$, $B_e = 3.3$, FC–FC$= 0.52 \pm 0.03$), we plot: **A** a heat map (carpet plot) of the full time-series simulation, and **B** the final 1 s of simulated dynamics for the six Medial brain regions (numbered 8–13).

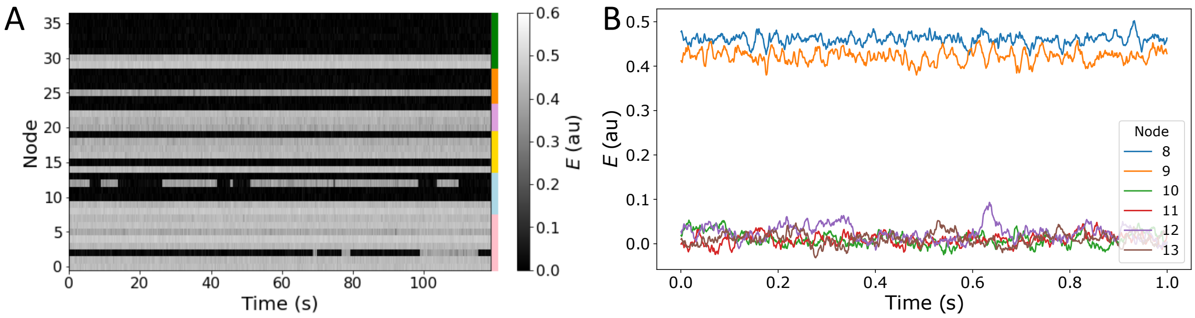


Figure S2: **Hysteresis simulation**. For the Hysteresis regime model with the best FC–FC fit ($G = 0.35$, $B_e = 3.7$, FC–FC$= 0.50 \pm 0.14$), we plot: **A** a heat map (carpet plot) of the full time-series simulation, and **B** the final 1 s of simulated dynamics for the six Medial brain regions (numbered 8–13). The carpet plot reveals evidence of long-timescale state switching for SSp-n (region 2) and RSPv (region 12).

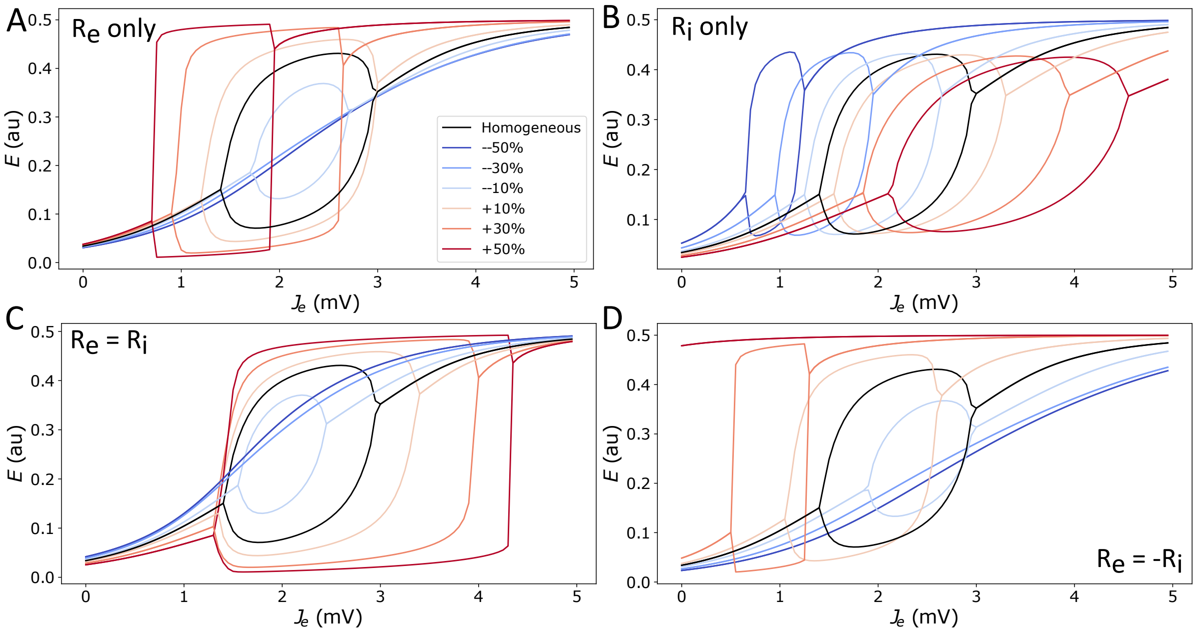


Figure S3: **The model's bifurcation structure in the Limit Cycle regime varies substantially when considering perturbations in local excitatory and inhibitory cell density, $R_e$ and $R_i$, of up to ±50%.** Instead of ±10% as in Fig. 5, here we plot perturbations of up to ±50%. These perturbations can have major effects on the bifurcation structure, including eliminating limit-cycle dynamics altogether.

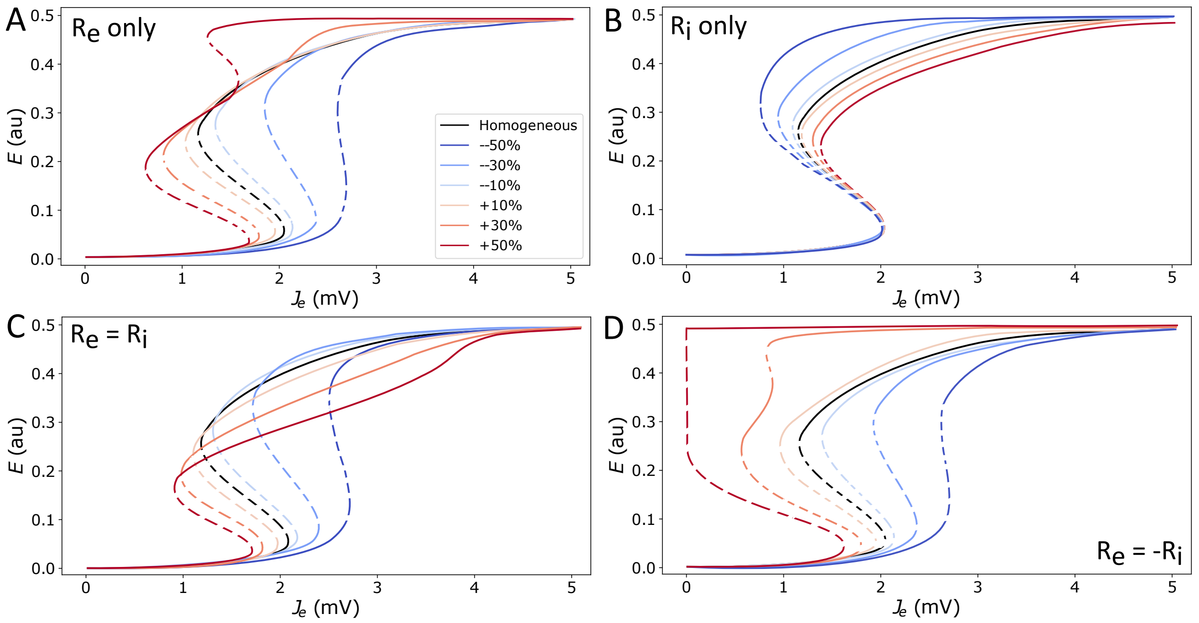


Figure S4: **The model's hysteresis bifurcation structure varies substantially when considering perturbations in local excitatory and inhibitory cell density, $R_e$ and $R_i$, of up to ±50%.** Of particular interest is the additional multi-stability via a new pair of saddle-node bifurcations (e.g., for $R_e = 0.5$).
